# A disease-causing Isoleucyl-tRNA synthetase variant leads to altered protein complex formation and cellular stress response

**DOI:** 10.1101/2025.08.21.671621

**Authors:** Han Gao, Felicia Pais Araújo, Jolie M. Miller, Samuel Protais Nyandwi, Juan Pablo Padilla-Martínez, Rasangi Tennakoon, Hui Peng, Haissi Cui

## Abstract

Aminoacyl-tRNA synthetases are key enzymes in protein synthesis, as they catalyze the attachment of amino acids to their designated, cognate tRNAs. As such, mutations in aminoacyl-tRNA synthetases are associated with severe diseases, such as neurodevelopmental disorders. Many of these mutations occur in the catalytically active site or tRNA binding domains, however, others can affect domains associated with multisynthetase complex formation.

Here, we investigate a disease-causing mutation in the UNE-L domain of Isoleucyl-tRNA synthetase (*IARS1*, IleRS), which mediates IleRS interactions within the multisynthetase complex. Interestingly, levels of the resulting protein were severely reduced in comparison to wildtype IleRS. While bulk protein synthesis and cell proliferation were not affected, the integrated stress response signaling pathway was altered. This change was exacerbated in low glucose medium, suggesting that mutant cells could respond differently to cellular stress. Our study hints at a possible underlying disease mechanism, where catalytic activity might not be affected but instead complex formation and protein stability.

## Introduction

Aminoacyl-tRNA synthetases (aaRSs) are a group of enzymes responsible for linking tRNAs with amino acids through esterification reactions to provide substrates for protein synthesis (1, 2). As such, they are essential for all life forms, as interpreters of the genetic code by recognizing both their corresponding amino acid and the cognate tRNA (3, 4). Charged tRNAs are then used by the ribosome for protein synthesis, where the sequence of amino acids in a protein is determined by base pairing of the tRNA anticodon to mRNA. AaRSs are named after the amino acid they recognize, for example, isoleucyl-tRNA synthetase (IleRS, IUPAC gene name *IARS1*) charges isoleucine on tRNA.

While aaRSs are generally active as either monomers or dimers, in multicellular organisms, certain aaRSs form a protein complex called the multisynthetase complex (MSC) (5). The composition of the MSC varies in terms of composition and complexity across species (6, 7). In mammals, the MSC includes eight aaRSs with nine enzymatic activities (ArgRS, GlnRS, MetRS, GluProRS, IleRS, LeuRS, AspRS, LysRS) (8). The MSC also contains three adaptor proteins, namely AIMP1, 2, and 3, which are essential for complex integrity (9–13). The individual proteins contain domains that facilitate the intermolecular interactions within the MSC (14), including leucine zippers (15) and Glutathione S-transferase-like motifs (16). Additionally, individual aaRSs can contain so called unique (UNE) domains, which can be involved in complex formation: The UNE-L domain in LeuRS (17) and the two UNE-I domains in IleRS mediate their integration into the MSC (18). Unlike other enzymes, where complex formation a prerequisite for function, aaRSs are enzymatically active both within and outside of the MSC (19). The function of the multisynthetase complex is therefore still ambiguous (20), as excluding individual aaRSs does not necessarily affect global protein synthesis (21), suggesting other functions through the regulation of non-canonical functions or in specialized translation.

In addition to their contributions to protein synthesis, aaRSs also mediate cell signaling through extensive expanded functions. These include but are not limited to the regulation of mRNA transcription (22) and translation (23–27) as well as cell signaling (28–30). Of note, LeuRS controls the mTOR pathway in response to glucose levels (30, 31) and IleRS is associated with BRCA and PI3K signaling (32, 33).

Due to the central role of aaRSs in protein synthesis, mutations can cause severe disorders in humans (34). These encompass disorders involving multiple organs, including the central nervous system, and often feature leukodystrophies (35). While many of these disease-causing mutations locate to the catalytic domain or affect tRNA binding, individual variants fall outside of domains classically associated with aaRSs’ enzymatic activity (36). The most common *RARS1* mutation in patients with hypomyelinating leukodystrophy leads to the preferential expression of a variant starting lacking the leucine zipper, which anchors ArgRS to the multisynthetase complex (37, 38). In a model cell line, decoding of arginine codons was not diminished and overall bulk protein synthesis as well as cell viability were comparable to control cells (21). Similarly, disease-associated point mutations have been found in the two UNE-I domains of IleRS (35), both of which are non-catalytic domains that mediate IleRS MSC integration and its interaction with LeuRS (14, 33, 39).

We were interested in exploring the mechanism underlying the pathogenicity of the IleRS variant c.3521T>A, I1174I>N (35), in the following referred to as I1174N, as interference with aminoacylation activity alone seemed unlikely due to its positioning in the UNE-I domain. We hypothesized that I1174N could alter the interaction of IleRS with other proteins, such as components of the multisynthetase complex. To understand the molecular consequences of the I1174N amino acid exchange in the UNE-I domain, we therefore aimed to develop a model cell line to explore how IleRS biochemistry and functions within the cell would be affected.

## Results

### Establishment of a c.3521T>A, p.I1174N IleRS model cell line

We used CRISPR/Cas9-based prime gene editing (40) to introduce the c.3521T>A mutation into a model cell line, HEK 293T, at the endogenous *IARS1* locus (Figure 1A). To this end, we designed guide RNA (gRNA), repair template, and second nick gRNA and inserted them into pEA1, a plasmid which carries the original Cas9-reverse transcriptase fusion protein (41). HEK 293T cells were then transfected with this plasmid and GFP-positive cells were sorted into individual wells (Supplementary Figure 1A). Following single cell selection, we retrieved ten colonies (Supplementary Figure 1B). The DNA sequence flanking the mutation site was amplified by PCR and the presence of the edit was verified by Sanger sequencing (Figure 1B, Supplementary Figure 1C). We identified five colonies which were heterozygous for the desired edit and four colonies which were homozygous (c.3521T>A, Figure 1B, Supplementary Figure 1C). Two clones with homozygous edits were selected for follow-up experiments; they will be referred to as I1174N_3 and I1174N_9 in the following.

**Figure 1:**
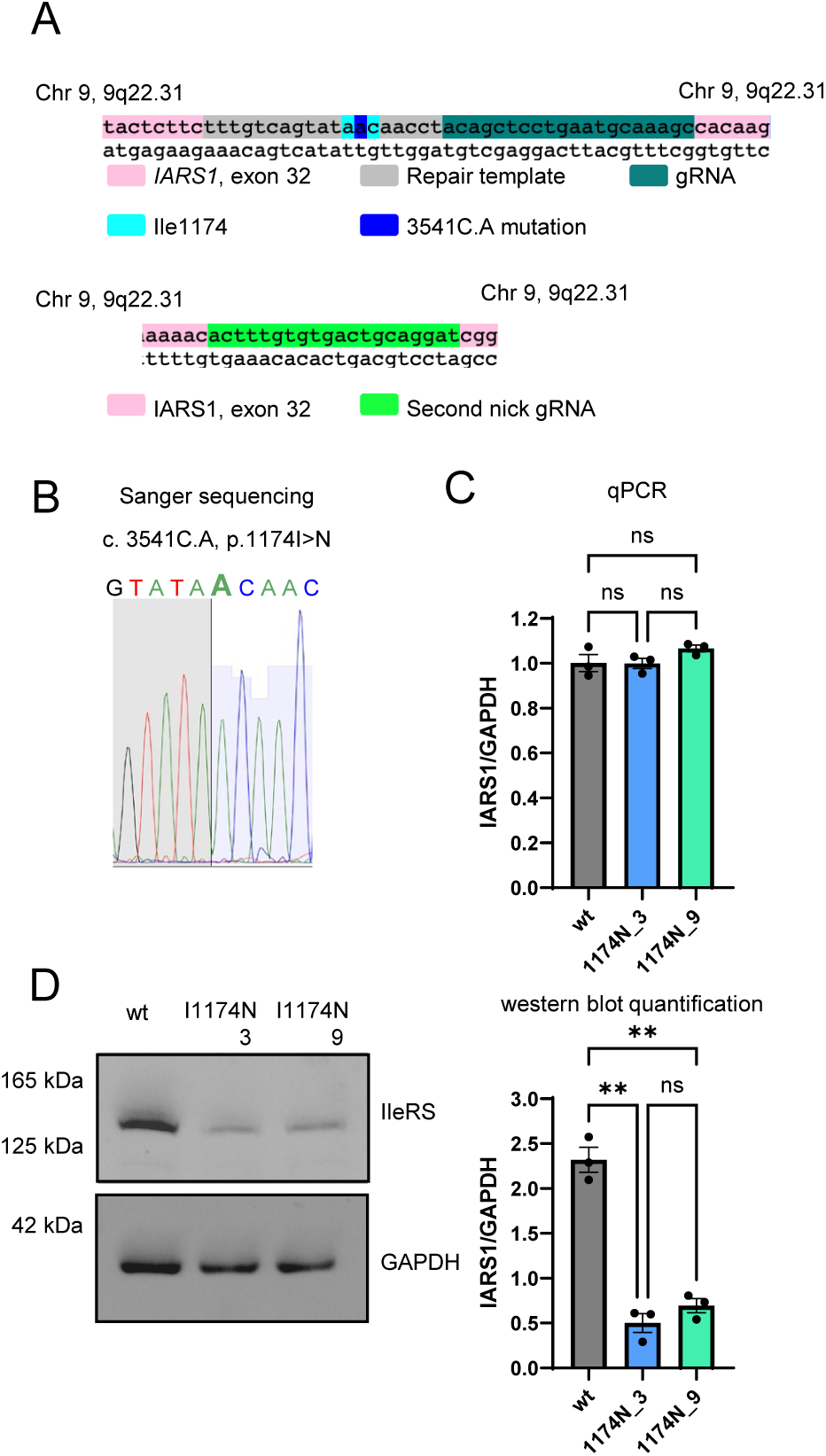
Characterization of a cell line model carrying a disease-causing mutations in IARS1. (A) Schematic representation of the c.3521T>A point mutation in *IARS1* exon 32, which leads to I1174N. The mutation involves a codon change from A**T**C (Isoleucine) to A**A**C (Asparagine), highlighted in dark blue within the cyan-shaded codon sequence. IARS1 exon 32 in pink, the repair template, gRNA, and second nick guide RNA in grey, dark green, and light green, respectively. (B) Sanger sequencing chromatogram of cells with a c.3521T>A point mutation in *IARS1* exon 32, which leads to I1174N. (C) RT-qPCR to determine IARS1 mRNA level in HEK293T WT and I1174N cells. IARS1 was normalized to the housekeeping gene GAPDH. (D) Representative western blot probing IleRS protein in HEK293T WT and I1174N cells and densitometric analysis. Results were normalized against GAPDH. (C, D) Data are presented as the mean of three biological replicates ± s.e.m. WT: wildtype HEK 293T cells. I1174N_3: IleRS I1174N variant clone 3. I1174N_9: IleRS I1174N variant clone 9.

### c.3521T>A mutation affects IleRS protein but not mRNA levels

After establishing the cell line, we investigated the impact of the mutation on IleRS mRNA and protein levels. To assess mRNA levels, we performed a two-step RT-qPCR using primers that span *IARS1* exons 26 and 27, positioned upstream of the mutation site. Primer efficiency was validated before the experiment, and GAPDH was used as a normalization control due to its stable expression across conditions. RT-qPCR results indicated no significant difference in *IARS1* mRNA levels between wildtype HEK 293T cells (WT) and c.3521T>A homozygous cells (Figure 1C).

However, upon assessing protein levels by immunoblotting, we found a 70% reduction in IleRS protein in I1174N homozygous cells compared to WT (Figure 1D). The molecular weight of IleRS did not change, suggesting that the amino acid change did not lead to premature termination (Figure 1D). Thus, while IleRS mRNA levels remained stable, the I1174N mutation significantly reduced protein levels.

### Global protein synthesis and cell viability were unaffected in the IleRS I1174N cells

Puromycin incorporation assays (42) were used to assess the impact of the IleRS I1174N variant (and in turn reduced IleRS protein levels) on global protein synthesis. Immunoblotting against puromycin and its densitometric quantification are shown in Figure 2A. Both WT and I1174N cell lines exhibited comparable signal intensities, suggesting that the I1174N exchange did not significantly affect the synthesis of new proteins. We further assessed cell viability and found that proliferation rates were similar between WT and IleRS I1174N cells, with no significant difference over 72 hours (Figure 2B). This suggests that the reduction of IleRS protein levels and therefore catalytic capacity *per se* was not limiting.

**Figure 2:**
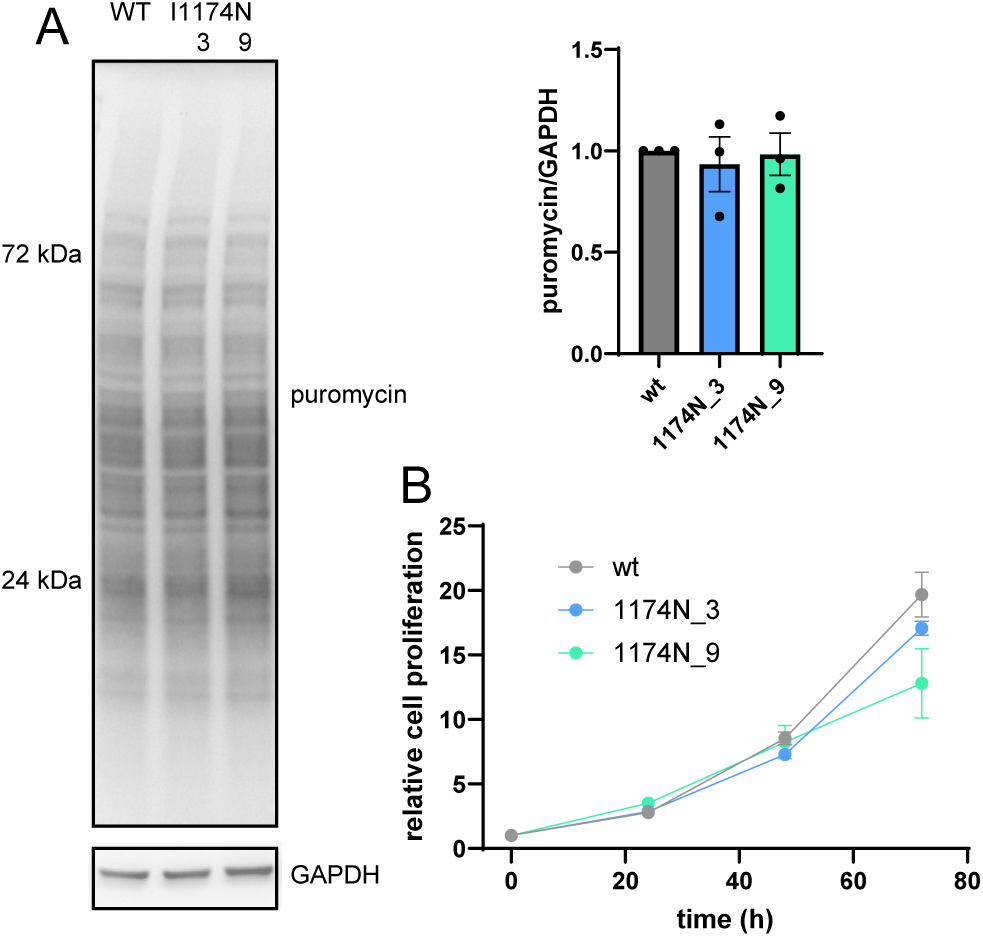
Bulk protein synthesis and cell viability were not affected by IleRS I1174N. (A) Left panel: Western blot to probe puromycin incorporation, which marks newly synthesized proteins. Right panel: Results were normalized against GAPDH. (B) Cell viability and cell proliferation on different days after seeding was assessed with Alamar Blue. (A, B) Data are presented as the mean of three biological replicates ± s.e.m. wt: wildtype 293T cells. I1174N_3: IleRS I1174N variant clone 3. I1174N_9: IleRS I1174N variant clone 9.

### I1174N mutation in the UNE-I domain of IleRS causes dissociation of IleRS from the multisynthetase complex

Since the UNE-I domain is crucial for the integration of IleRS into the MSC (39, 43), we hypothesized that the I1174N exchange, which falls in the UNE-I domain, might disrupt the association of IleRS with other proteins in the complex. To investigate the overall integrity of the MSC as well as the presence of IleRS, we performed size exclusion chromatography. Under mild cell lysis conditions, the MSC stays intact and proteins in fractions corresponding to different molecular weights can be detected through immunoblotting. We probed MSC-bound aaRSs (IleRS, ArgRS, LeuRS) and non-MSC-bound TyrRS to assess whether IleRS was still bound in the MSC and would therefore elute at earlier fractions. Immunoblotting of size exclusion chromatography fractions of WT cell lysates (Figure 3A) showed a prominent IleRS signal at fraction 9-10 in WT cells, in line with MSC association. In contrast, the I1174N cell line displayed strongly reduced signal intensity at fraction 9-10 in line with lower overall levels in the cell, confirming reduced IleRS presence in the MSC (Figure 3A). Shorter IleRS variants eluting at fraction 11, which were also present in WT, in turn constitute a larger percentage of total IleRS protein in I1174N cells (Figure 3A).

**Figure 3:**
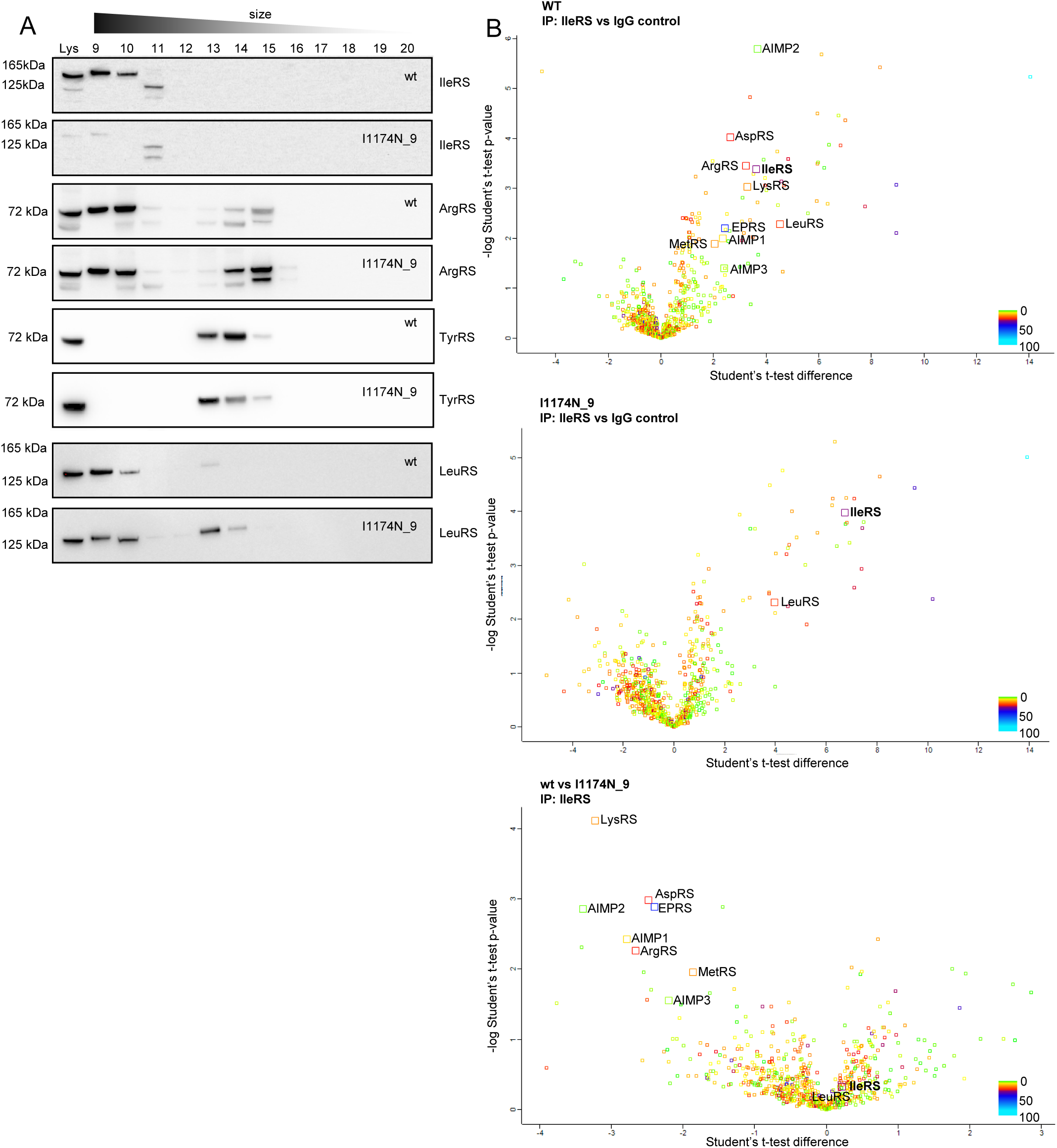
IleRS I1174N caused the exclusion of IleRS and LeuRS from the multisynthetase complex. (A) Cell lysate separated by size exclusion chromatography to distinguish between monomeric/dimeric and multisynthetase complex-bound aminoacyl-tRNA synthetases (aaRS) followed by western blot. Intact multisynthetase complex elutes between fractions 9–11. Dimeric or monomeric tRNA synthetases elude between fractions 13–15. IleRS: Isoleucyl-tRNA Synthetase. ArgRS: Arginyl-tRNA Synthetase. TyrRS: Tyrosyl-tRNA Synthetase. LeuRS: Leucyl-tRNA Synthetase. Data is shown as a representative of at least three biological replicates. (B) Volcano plot of proteins identified by mass spectrometry following co-immunoprecipitation. Color coding denotes the number of peptides per protein. (A, B) wt: wildtype HEK 293T cells. I1174N_9: IleRS1 I1174N variant clone 9.

We next explored the influence of IleRS I1174N on other MSC subcomplexes. ArgRS immunoblotting showed no major effect of the IleRS mutation on ArgRS MSC association, with strong bands at fractions 9-10 in both WT and mutant cells, suggesting that the majority of the MSC is still intact (Figure 3A). However, ArgRS bands at lower molecular weight fractions increased which could indicate that complexes might be less stable compared to WT cells (Figure 3A). TyrRS eluted consistently between fractions 13-15 in both cell lines, as expected for a non-MSC-bound protein (Figure 3A).

LeuRS is tethered to the MSC through IleRS and we hypothesized that LeuRS would also dissociate if IleRS binding to the MSC is disrupted. In WT cells, full-length LeuRS was mostly detected in fractions 9 and 10 (Figure 3A). However, in I1174N cells, LeuRS appeared in fractions 9-10 and in comparable intensity in fractions 13 and 14, corresponding to free LeuRS (Figure 3A). This indicates that the reduced integration of IleRS into MSC in turn affected LeuRS.

We further verified the loss of multisynthetase-complex defining interactions in IleRS I1174N cells through interactome studies. We therefore co-immunoprecipitated IleRS from both WT and I1174N cells and identified interaction partners through mass spectrometry (Figure 3B, Supplementary Table S1, S2). Pulldown of IleRS and enrichment of its known interaction partner LeuRS was verified with western blot (Supplementary Figure 2A). Given the unexpectedly large number of interactors, we opted to repeat the co-immunoprecipitation with a second polyclonal antibody to control for possible antibody-off target effects (Supplementary Figure 2B, C, Supplementary Table S2). While in both experiments additional interactors were found, only members of the MSC were present in both (Supplementary Figure 2D).

In WT cells, the IleRS interactome included all MSC components, except GlnRS, for which insufficient peptides were detected (Figure 3B). GlnRS was present in pulldowns with the second IleRS antibody (Supplementary Figure 2C). In contrast, in I1174N cells, IleRS enrichment only retrieved LeuRS, while all other MSC components were absent (Figure 3B). Using the alternative antibody, IleRS enrichment was reduced in I1174N cells and the LeuRS interaction was also reduced along with other MSC components (Supplementary Figure 2C), which could either be due to a difference in binding site or reflective of lower IleRS enrichment in this experiment. This confirms our size-exclusion chromatography results, suggesting again that the I1174N exchange disrupts IleRS association with MSC proteins.

### IleRS I1174N variant affects levels of the stress response protein ATF4

In order to understand how the I1174N variant might affect cellular processes, we tested several cells signaling pathways that were previously associated with aaRSs variants and disease progression: Mutations in aaRSs are known to cause changes in the integrated stress response, most commonly its increased activation (44–46). As the integrated stress response reports on uncharged tRNAs and perturbations in amino acid homeostasis (47), it is specifically vulnerable to alterations in aminoacylation. We therefore probed for increases in Activating Transcription Factor 4 (ATF4) protein levels, as ATF4 is a central regulator of the integrated stress response (48, 49). Immunoblotting was conducted on cells subjected to different cell stress conditions: low glucose, hypoxia mimicry, and serum starvation (50) (Figure 4A). Under serum starvation and hypoxia mimicry, ATF4 levels were upregulated compared to control conditions, but the ATF4 response was similar between WT and I1174N cells. In contrast, in low glucose medium, ATF4 levels in I1174N cells were lower than those in WT cells (Figure 4A). Total protein synthesis, visualized by puromycin incorporation, was unchanged, suggesting that at altered ATF4 level did not impact bulk protein synthesis (Figure 4B).

**Figure 4.**
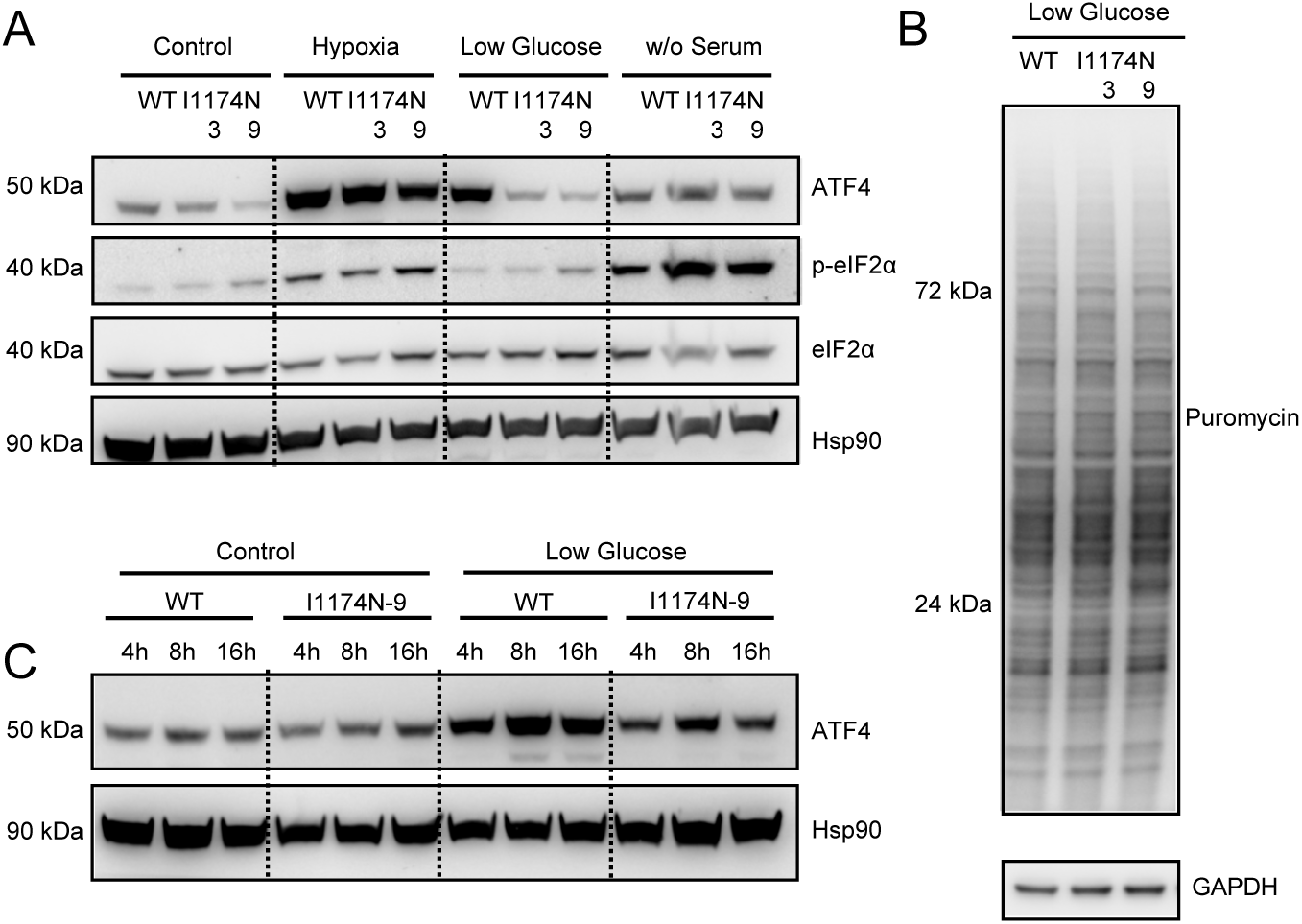
Reduced induction of integrated stress response regulator ATF4 in low glucose medium in IARS1 I1174N cells. (A) Protein levels of ATF4, phosphorylated eIF2α (p-eIF2α), total eIF2α, and Hsp90 were analyzed by western blot under various cell stress conditions. Hypoxia: Hypoxia-mimicry through addition of iron chelator DFO. Low Glucose: Low glucose medium. W/O serum: Serum starvation through removal of FBS. Hsp90 was used as a loading control. Data shown is representative of at least three biological replicates. (B) Western blot to probe puromycin incorporation in low glucose medium, which marks newly synthesized proteins. (C) ATF4 levels over time in low glucose medium. (A-C) WT: wildtype HEK 293T cells. . I1174N_3: IleRS I1174N variant clone 3. I1174N_9: IleRS I1174N variant clone 9.

To investigate how fast a difference in ATF4 regulation would be detectable, we tested ATF4 levels at different time points after initiating reduced glucose levels. At basal, high glucose levels, ATF4 protein was comparable between WT and I1174N cells (Figure 4C). At 4 h, 8 h, and 16 h after glucose starvation, a reduction in ATF4 levels was observed in cells with the I1174N IleRS variant (Figure 4C). To test whether ATF4 target gene expression was changed, we used qPCR to assess levels of CHOP, HSPA5, EIF2S2, EPRS, YARS, and SARS mRNA after 24 hours in low glucose medium. While we would expect downregulation of ATF4 target genes, we did not find consistent changes (Supplementary Figure 3A, B).

The mTOR pathway controls the cellular response to nutrients such as amino acids. As LeuRS controls mTOR signaling in response to glucose fluctuation (31, 51), we also assessed the mTOR pathway in I1174N cells by quantifying levels of its downstream targets p-S6K1 and p-4EBP1 (Supplementary Figure 4A, B). p-4EBP1 levels were unchanged while p-S6K1 levels were subtly increasing in I1174N cells (Supplementary Figure 4A). Levels of phosphorylated ribosomal protein S6 kinase 1 (p-S6K1) on the other hand dropped over time in WT cells and low glucose medium exacerbated this effect (Supplementary Figure 4B). I1174N cells, however, showed increased p-S6K1 levels after 8 h in comparison to WT due to a slower decrease over time (Supplementary Figure 4B), further suggesting that the cellular response to low glucose levels is attenuated in I1174N cells.

## Discussion

Here, we evaluate the impact of a disease-causing variant, IleRS I1174N, with regard to its association with other multisynthetase complex proteins, consequences on cell growth and protein synthesis, the IleRS interactome, and cellular stress signaling. We found that the I1174N amino acid exchange strongly decreased IleRS protein levels while mRNA levels remained constant. The resulting reduction of IleRS did not negatively affect bulk protein synthesis or cell viability. The I1174N amino acid exchange in the UNE-I domain disrupted interactions between IleRS and other MSC-bound aaRS and increased the amount of free LeuRS. Levels of the integrated stress response regulator ATF4 were reduced in low glucose conditions.

In the affected individual (35), I1174N is accompanied by another mutation on the other allele, c.1252C.T,p.418R>*, which truncates IleRS within its catalytic domain and deletes the IleRS tRNA binding domain, very likely resulting in catalytically inactive protein. Neither variant by itself was disease causing, as evident by the unaffected parents and siblings (35), suggesting that the activity resulting from one allele is sufficient to support aminoacylation needs. However, the effects of I1174N and R418* seem to be additive and together lower IleRS functionality to a critical degree. Homozygous cases with I1174N have not been reported and it therefore difficult to predict whether I1174N alone would be sufficient to elicit a disease phenotype. We opted to explore a homozygous cell line model to allow focused and unambiguous studies on the biochemical and cell biology consequences of the I1174N variant.

The mammalian MSC can be divided into three subcomplexes (14): Subcomplex III is a trimeric complex formed by EPRS, IleRS, and LeuRS. Within this structure, IleRS interacts with EPRS via its UNE-I domain and anchors LeuRS, which binds to the catalytic domain of IleRS through its UNE-L domain. EPRS then facilitates the connection between subcomplexes II and III. In line with this model, the I1174N amino acid exchange in the UNE-I domain seems sufficient to disrupt the interactions between IleRS and EPRS, without affecting leRS and LeuRS interaction, as LeuRS was still enriched upon pulldown of the IleRS I1174N variant while interaction with EPRS and all other MSC-bound proteins was lost (Figure 3B). Interestingly, in the co-crystal structure of the chicken EPRS-IleRS complex (PDB 7WRS) (33), I1174 is not directly in contact with EPRS and the closest contact to EPRS is over 10 Å removed. Alphafold places amino acid 1174 in a beta sheet in the middle of the human UNE-I domain, and the exchange to Asn is not predicted to disrupt its fold. As EPRS binds in a cleft between the two UNE-I domains (33), the I1174N exchange might however disrupt the careful spacing between the two or otherwise indirectly interfere with protein-protein interactions.

The integration of IleRS into the multisynthetase complex can influence protein functionality but also stability: A previous study has shown that the levels of aaRSs in the MSC can be dependent on each other, as knock down of individual components led to a depletion of others (43). Reduction of the adapter proteins AIMP-2 and AIMP1 as well as EPRS, caused IleRS levels to decrease (43), suggesting that IleRS outside of the MSC is less stable. In turn, IleRS knockdown was also shown lower levels of MetRS and LeuRS (43). We confirmed that the reduction in IleRS also led to a reduction in LeuRS protein with downstream consequences for mTOR signaling (Supplementary Figure 4A), allowing for IleRS to impact a LeuRS-mediated signaling pathways through the multisynthetase complex.

Previous studies on the role of the multisynthetase complex suggest that its functions might be multifaceted: A channeling effect of the MSC, meaning the increased the flow of charged tRNAs to the ribosome, has been discussed (52). Specific genes are differentially translated upon exclusion of ArgRS and GlnRS from the MSC (21), suggesting that the contribution of the MSC to mRNA translation could affect protein synthesis differently across transcripts. However, bulk translation seems to be less sensitive to the disruption of MSC formation (53). We found that bulk protein synthesis was similarly unaffected by the reduction of IleRS levels and its altered MSC-association (Figure 2A). In line with this, cell viability was unaltered in I1174N cells, and proliferation rates were comparable to WT cells (Figure 2B). Cell stress through low glucose did not exacerbate this phenotype as protein synthesis was likely still sufficient.

The integrated stress response mediates the cellular adaptation to different environmental triggers, such as dsRNA exposure, signifying viral infection, accumulation of unfolded proteins, and amino acid starvation (47). Over-activation of the integrated stress response is a strong contributor to aaRS-mutation induced Charcot-Marie-Tooth disease, a neurodegenerative disorder which predominantly affects long peripheral motor neurons (54). In *GARS1* CMT mutations, sequestration of tRNAs by disease-causing GlyRS variants induced an overactivation of the integrated stress response (44–46). Here, we found the opposite, where a disease-causing variant of IleRS led to a reduced activation of ATF4 (Figure 4). A recent study showed that inhibition of aaRSs enzymatic activity prevented RNP granule assembly upon exogenous stress (55), offering a possible pathway as to how IleRS I1174N could attenuate the integrated stress response.

In summary, we found the marked reduction of IleRS protein in a cell line modelling a disease-causing variant together with the exclusion of IleRS from its native protein complex. These observations went hand in hand with an impaired induction of the integrated stress response in a nutrient low environment. This suggests that reduced cellular adaptation to stress could be a potential factor in IleRS-driven disease elicited by mutations in *IARS1*.

## Material and Methods

### Cell culture

HEK 293T cells were obtained from ATCC and confirmed by STR analysis. Cells were cultured at a density of 80-90% confluency and passaged on average twice per week. If not stated otherwise, cells were maintained in high glucose DMEM (Sigma-Aldrich), with 10% FBS (Corning), and 1% Penicillin/Streptomycin (Sigma-Aldrich). Cells were tested monthly for mycoplasma contamination and were consistently found negative. Experiments were performed before passage 30.

### Prime editing and single cell sorting

The PEA1-GFP plasmids used here were generated with guide RNA (gRNA), repair template, and second nick gRNA designed using Petal (41). The plasmid was transfected into HEK293T cells using Calfectin (SignaGen). Successfully transfected cells were selected based on GFP expression by the Temerty Faculty of Medicine flow cytometry facility and sorted as individually into a 96-well plate.

### Sanger sequencing

DNA was isolated with DNA extract solution containing proteinase K (QuickExtract DNA Extraction Solution, Lucigen). Primers were designed to amplify 596 bp surrounding the c.3521T>A site in a two-step PCR using Phusion polymerase (NEB). The molecular weight of the amplified product was confirmed by 1% agarose gel electrophoresis. The product was then purified with a PCR purification kit (FroggaBio) and 300 ng of DNA were sequenced with a custom primer at the The Centre for Applied Genomics, SickKids, Toronto.

### Protein extraction

Cells were seeded at a density of 5 × 10^6^ cells into 10 cm dishes one day prior to harvest. After washing with PBS (Sigma Aldrich) twice, total protein was collected in 500 μl RIPA lysis buffer (0.05 M Tris/HCl, pH = 7.4, 0.15 M NaCl, 0.25% deoxycholate, 1% NP-40, 1 mM EDTA) with protease inhibitor cocktail (cOmplete™, Mini, EDTA-free Protease Inhibitor Cocktail, Roche). Cells underwent freeze-thaw cycles to ensure complete lysis and were stored at -80°C. Lysates were centrifuged at 14,000 x g for 10 minutes and the pellet was removed. SDS-loading buffer (5x: 0.25% Bromophenol blue, 0.5 M dithiothreitol, 50% glycerol, 10% sodium dodecyl sulfate, 0.25 M Tris-HCL) was added and the lysates were heated at 95 degrees for 7 minutes prior to SDS PAGE.

### Western blot

3-5 μl (6-10 μg protein) of lysates were loaded on 4-12% gradient gel for gel electrophoresis. Transfer to PVDF membranes was performed with the Invitrogen iBlot 2 Dry Blotting system. Membranes were blocked in 3% milk/TBST for 1 hour. After washing with TBST, membranes were incubated with the respective primary antibody in 5% BSA/TBST overnight at 4 degrees. The next day, membranes were washed 3 times for at least 10 minutes with TBST. Membranes were then incubated with the corresponding HRP-conjugated secondary antibody diluted in 1% milk/TBST (Goat-anti-rabbit, Invitrogen, 1:5000, Goat-anti-mouse, Invitrogen, 1:10000) for 60-90 minutes and washed. Western blots were visualized with chemiluminescence (SignalFire™ ECL Reagent, Cell signaling) and imaged with a ChemiDoc (Bio-Rad). All antibodies used here are listed in the Supplementary Table S3.

### RNA isolation

Cells were seeded into 10 cm dishes two days prior to the harvest at a density of 3 × 10^6^ cells/dish. After washing with PBS (Sigma Aldrich) twice, RNA was extracted with TRIzol (Invitrogen) following the manufacturer’s instructions. RNA integrity was confirmed 1% agarose gel electrophoresis.

### RT-qPCR

cDNA was synthesized from extracted RNA using the High-Capacity cDNA Reverse Transcription Kit (Invitrogen) according to the manufacturer’s instructions. A LightCycler® 480 System (Roche) was used to analyze cDNA levels and values were normalized to GAPDH/18S. 2^-ΔΔCt values are shown. All the qPCR primers used here can be found in Supplementary Table S4.

### Size exclusion chromatography to assess protein complex formation and integrity

Cells were seeded into 15 cm dishes one day prior to the experiment at a density of 5 x 10^6^ cells/dish. After washing with PBS (Gibco) twice, cells were harvested by adding 500 μl MSC lysis buffer (20 mM Tris–HCl, pH 7.5, 1 mM EDTA, 150 mM NaCl, 1% NP-40 and Protease Inhibitor Cocktail cOmplete™, Mini, EDTA-free Protease Inhibitor Cocktail, Roche). The lysates were incubated on ice for 20 minutes, followed by a 10-minute spin at 14000 x g to obtain soluble proteins. 500 μl of cell lysates were then injected onto a Superdex 200 Increase Tricon™ 10/300 GL Prepacked SEC Column (Cytiva) equilibrated with PBS (Gibco) and 1 ml fractions were collected. 20 μl of each fraction was analyzed by western blot.

### Puromycin incorporation assay

Assays were performed as described in (42). In brief, cells were seeded into a six-well plate one day prior at a density of 5 × 10^5^ cells/well. Puromycin was directly added at 10 μg/ml for 30 minutes (pulse). The medium was then changed (chase), and cells were allowed to recover for an additional one hour before lysis in RIPA buffer and analysis by western blot.

### Cell proliferation assay

Cells were seeded in quadruplets in 96-well cell culture plates at 5000 cells/well. One plate was used for every time point. 10% Alamar blue reagent (Invitrogen) were added 2 hours prior to fluorescence intensity measurements (λex = 570 nm, λem = 600 nm) with a microplate reader (Synergy H1, Biotek).

### Co-immunoprecipitation

293T WT or I1174N mutant cells were seeded onto 15 cm dishes at a density of 15 × 10^6 cells per dish, one day prior to lysis. Cells were harvested in 1 mL of mild lysis buffer (Tris-buffered saline with 1% NP-40 and 1 mM MgCl_2_). Cells were incubated on ice for 30 minutes, followed by centrifugation at 14,000 × g for 20 minutes to remove insoluble components. 1 ml supernatant was loaded onto 30 µl of pre-equilibrated protein A/G agarose beads (SCBT) along with 3 μL anti-IleRS antibody (Proteintech or Bethy Laboratories). After 3 hours, beads were washed once with 1 ml 0.05% NP-40 in TBS and twice with 1 ml TBS. The beads were then stored in PBS (Gibco) at -80° C.

### Protein digestion and Mass spectrometry

Mass spectrometry experiments were performed as described in (56). Peptides were eluted from protein A/G agarose beads by trypsin digest (5 ng/µL trypsin, 2 M urea, 50 mM Tris-HCl, pH 7.5, 0.2 mM DTT, 1 mM chloroacetamide). The digested peptides were desalted with C18 pipette tips (Pierce) according to the manufacturer’s instructions. Peptides were eluted in a final volume of 30 µL, concentrated using a SpeedVac (Thermo Fisher), and stored at -80°C. Prior to analysis, peptides were resuspended in 40 µL of 1% formic acid.

Samples were analyzed using an EASY-nLC 1200 UHPLC system combined with a Q-Exactive HF-X orbitrap mass spectrometer via an in-line nanoLC-electrospray ion source (Thermo Fisher Scientific). Solubilized peptides were loaded onto a house-made fused- capillary silica precolumn (100 μm I.D.) packed with 2 cm of 5 μm Luna C18 100 Å reverse phase particles (Phenomenex) and separated on a 14.1 cm (100 μm I.D) silica pulled emitter packed with 1.9 μm Luna C18 100 Å reverse phase particles (Phenomenex); capillary was sourced from Polymicron Technologies. Peptides were eluted over 120 min with mobile phase A (0.1% FA) and mobile phase B (80/20/0.1 ACN/water/FA) at a constant flow rate of 300 nL/min with a linear increase from 2% to 5% B over 1 min, an increase to 26% B over 70 min, an increase to 60% B over 20 min, then by an increase to 100% B over 14 min, and a final plateau at 100% B for 15 min.

Mass spectrometry was performed, as described by Hall, Yeung, and Peng (57). In brief, a data-dependent top 20 method was used. Full MS scans were acquired from *m/z* 350– 1400 at a resolution of 60,000 at 200 *m/z* with a target automated gain control (AGC) of 3 × 10^6^ charges. For higher-energy collisional-dissociation MS/MS scans, the normalized collision energy was set to 28 and a resolution of 15,000 at 200 *m/z*. Precursor ions were isolated in a 1.4 *m/z* window and accumulated for a maximum of 20 ms or until the AGC target of 1 × 10^5^ ions was reached. Precursors with unassigned charge states, a charge of 1+, or a charge of 7+ and higher were excluded from sequencing. Previously targeted precursors were dynamically excluded from re-sequencing for 20 s.

Mass spectrometry data were processed with MaxQuant 2.6.4.0 and Perseus 2.1.2 (58, 59) as described previously (56) to identify IleRS interaction partners. In brief, raw files were searched against Homo sapiens reference proteome UP000005640_9606 (60) (Uniprot) with the Andromeda search engine integrated into MaxQuant. Default settings were used for LFQ. The resulting protein groups file was loaded into Perseus and filtered for ‘reverse’, ‘potential contaminants’ and ‘only identified by site’. The log2 values of LFQ intensities were calculated, and all proteins with less than two valid values/group were discarded. Missing values were replaced from a normal distribution (width 0.3 and downshift 1.8), and Students t-test was used to calculate t-test significance and difference.

### Induction of cell stress

Cells were seeded into a six-well plate at a density of 1 × 10^6^ cells per well, one day prior to treatment. Cells were washed with PBS (Gibco) before introducing fresh medium. The control group was maintained in high glucose DMEM (Sigma-Aldrich) supplemented with 10% FBS (Corning) and 1% Penicillin/Streptomycin (Sigma-Aldrich). To mimic hypoxia group, 100 μM deferoxamine (Sigma-Aldrich) was added. Cells in serum-free conditions were exposed to DMEM without FBS and low glucose DMEM (Sigma-Aldrich) with 10% FBS and 1% Penicillin/Streptomycin was used for low glucose conditions. Cells were exposed for stressors for the indicated time and then harvested for western blot analysis.

### Statistics

Statistical significance was evaluated by comparing biological replicates with either Student’s t-test or ANOVA, respectively (Prism GraphPad). Mass spectrometry data was analyzed using Perseus (59).

## Acknowledgements

We thank the Flow cytometry facility at the University of Toronto for assistance with cell sorting. Further, we thank the The Centre for Applied Genomics at The Hospital for Sick Children for Sanger sequencing as well as the Donnelly Sequencing Centre for tRNA sequencing. HC acknowledges funding by the Canadian Institutes of Health Research (CIHR) PJT 497271, CFI/JELF, the Department of Chemistry, and the Faculty of Arts and Science at the University of Toronto. FPA and JPPM were supported by a MITACS Globalink Research Internship. SPN was supported by CIHR Canada Graduate Scholarships – Master’s and Doctoral program.

## Author contributions

HC designed research. HG, FPA, JMM, SPN, JPPM, and RT performed research. HG, FPA, JMM, and HC analyzed data. HC and HP supervised research. HC and HG wrote the manuscript with input from all authors.

## Supplementary Information

**Supplementary Figure 1.**
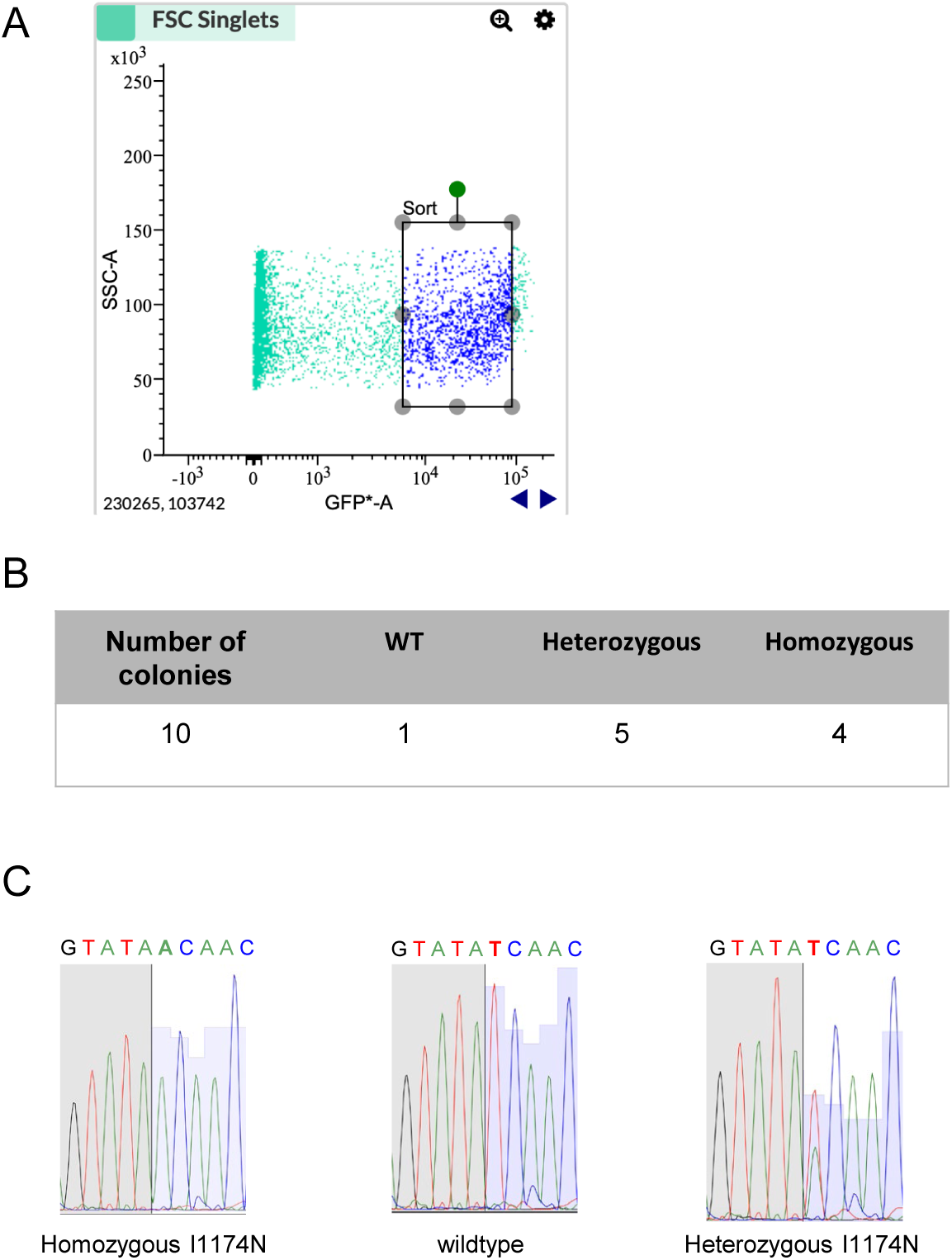
Establishment of an IleRS I1174N cell line. (A) Fluorescence activated cell sorting to identify successfully transfected cells. (B) Distribution of genotypes in monoclonal cell lines. WT: wildtype HEK 293T cells. (C) Representative Sanger sequencing results of *IARS1* regions flanking c.3521T>A.

**Supplementary Figure 2.**
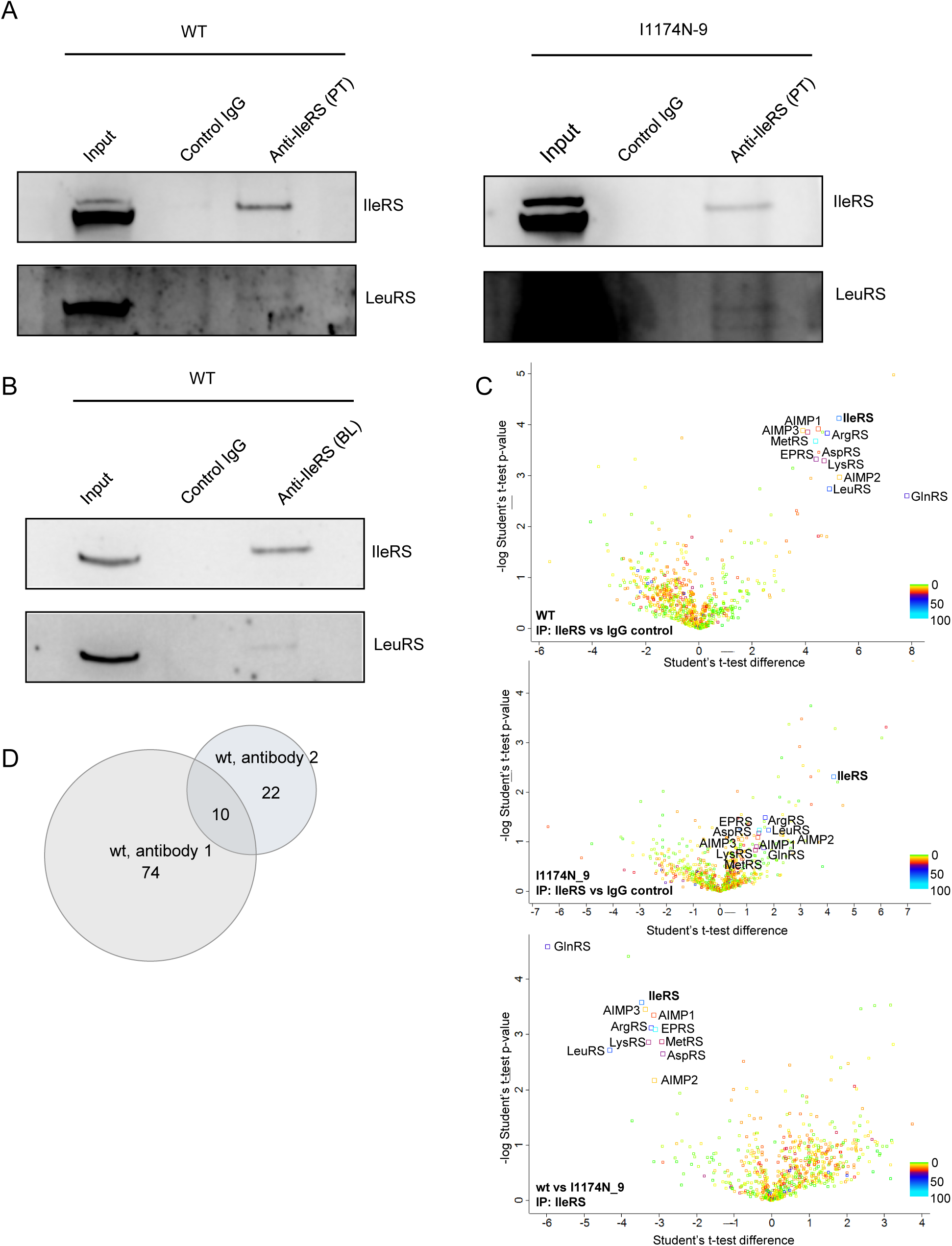
Verification of co-immunoprecipitation and interactome study. (A) Co-immunoprecipitation (Co-IP) and western blot following enrichment of IleRS. The IleRS interactor LeuRS was probed to confirm Co-IP. (B) Co-immunoprecipitation (Co-IP) and western blot following enrichment of IleRS with a second polyclonal antibody. The IleRS interactor LeuRS was probed to confirm Co-IP. (C) Interactome of IleRS in wt and mutant cell lines. (D) Comparison of interactomes obtained with two different IleRS antibodies. (A, C) WT: wildtype HEK 293T cells. I1174N_9: IleRS I1174N variant clone 9.

**Supplementary Figure 3.**
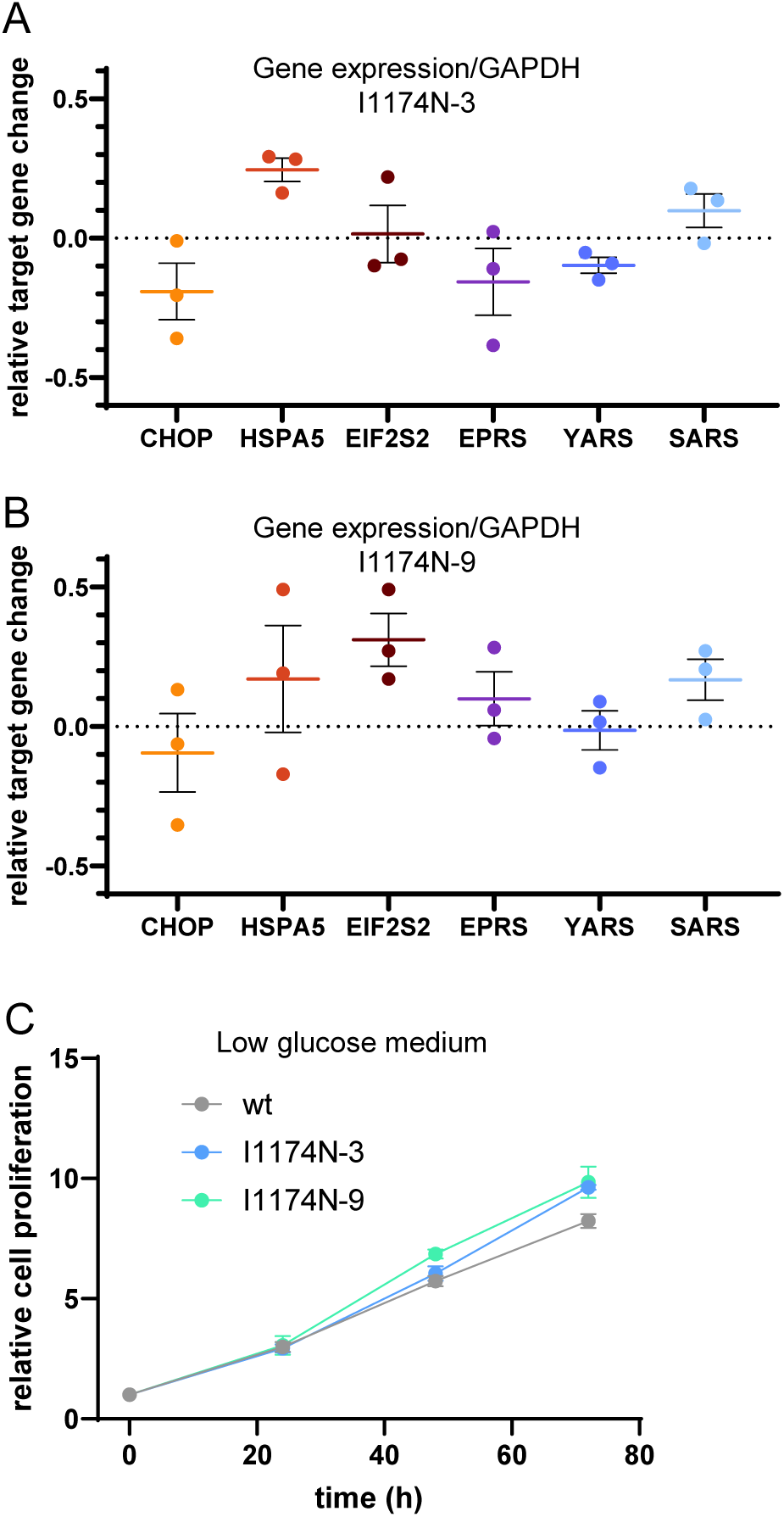
(A) qPCR of ATF4 target genes shows inconsistent regulation in IARS I1175N cells after 24 hours in low glucose medium. qPCR is shown relative to wildtype cells. (B) Cell proliferation in low glucose medium. (A, B) Three biological replicates ± s.e.m are shown. WT: wildtype HEK 293T cells. I1174N_3: IleRS I1174N variant clone 3. I1174N_9: IleRS1 I1174N variant clone 9.

**Supplementary Figure 4.**
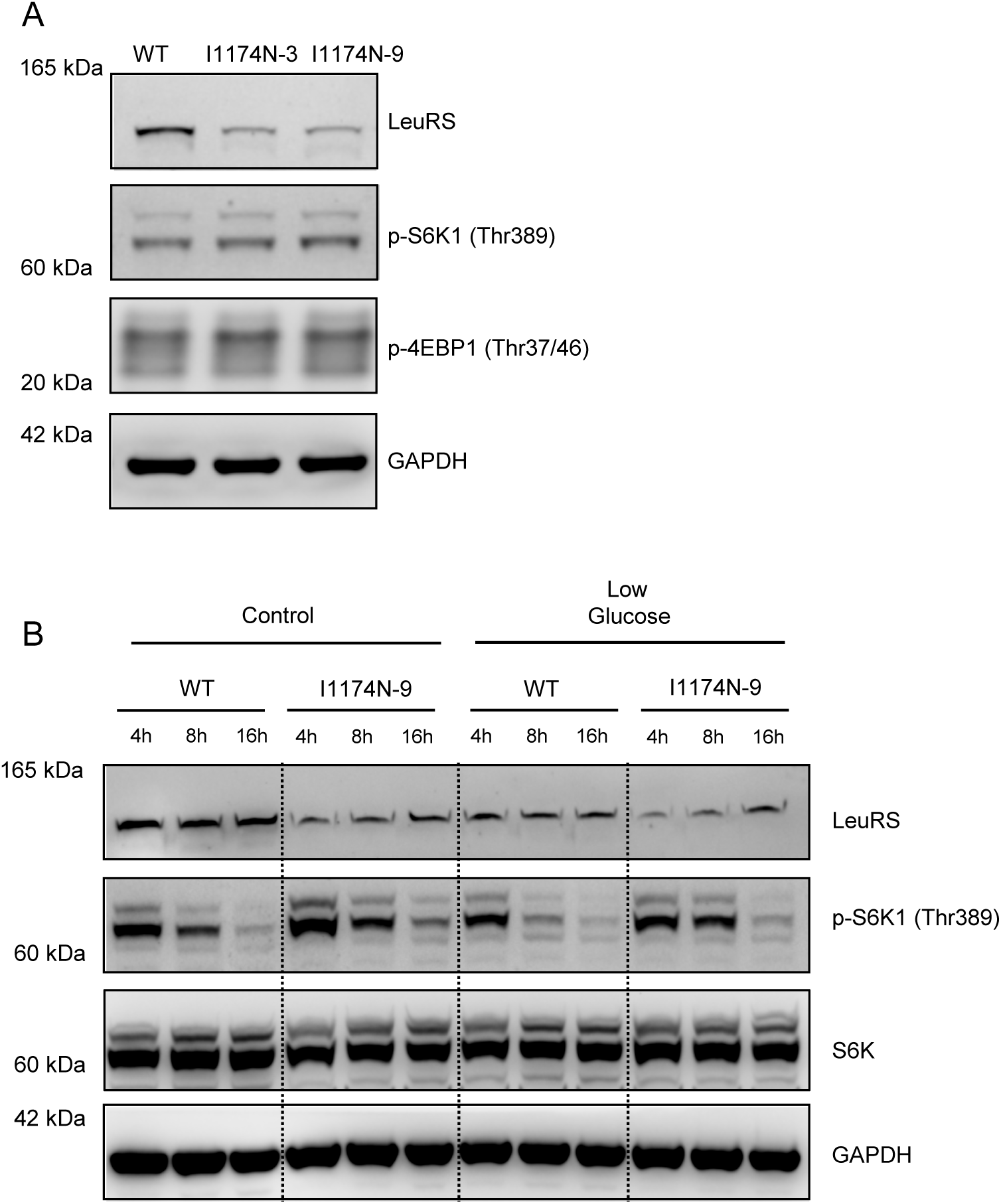
(A) mTOR signaling in I1174N cells as indicated by the phosphorylation status of S6K1. (B) mTOR signaling in I1174N cells in low glucose medium as indicated by the phosphorylation status of S6K1. (A, B) WT: wildtype 293T cells. I1174N_9: IleRS1 I1174N variant clone 9.

